# Cas9-cleavage sequences in minimal plasmids enhance non-viral genome targeting of CARs in primary human T cells

**DOI:** 10.1101/2020.12.31.424920

**Authors:** Ruirui Jing, Peng Jiao, Jiangqing Chen, Xianhui Meng, Xiaoyan Wu, Yanting Duan, Kai Shang, Liling Qian, Yanjie Huang, Junwei Liu, Tao Huang, Jin Jin, Wei Chen, Xun Zeng, Weiwei Yin, Xiaofei Gao, Chun Zhou, Michel Sadelain, Jie Sun

**Author notes:** These authors contributed equally to this work (Ruirui Jing, Peng Jiao).

## Abstract

T cell genome editing holds great promise to advance a range of immunotherapies but is encumbered by the dependence on difficult-to-produce and expensive viral vectors. Here we have designed small double-stranded plasmid DNA modified to mediate high-efficiency homologous recombination. The resulting chimeric antigen receptor (CAR)-T cells display a similar phenotype, transcriptional profile and *in vivo* potency as CAR-T cells generated using adeno-associated viral (AAV) vector. This method should simplify and accelerate the use of precision engineering to produce edited T cells for research and clinical purposes.

---

The US FDA has approved three CD19-specific chimeric antigen receptor (CAR)-T products to treat several hematological malignancies. These products use either retroviral (RV) or lentiviral (LV) vectors to deliver CAR genes into primary human T cells^1^. Both RV and LV vectors integrate in semi-random fashion, resulting in variegated transgene expression and the risk of insertional mutagenesis^2,3^. In contrast, genome editing can precisely insert genes at defined genomic sites using homology-directed repair (HDR)^4–7^. Using CRISPR/Cas9 to insert a CD19-specific CAR at the T-cell receptor α constant (*TRAC*) locus will at once delete expression of the endogenous T cell receptor (TCR) and afford effective CAR expression, resulting in reduced T cell exhaustion and improved tumor rejection in a mouse model of leukemia^8^. Expression and regulation of *TRAC*-encoded CAR reduces tonic signaling, delays differentiation and decreases exhaustion of T cells, enhancing potency of eradicating xenograft tumor^8^. Targeting different CARs as well as TCRs to *TRAC* locus showed similar enhancement compared to their virus-transduced counterparts^9–12^. Genome edited T cells may also be safer as TCR ablation minimizes the risk of autoimmunity and alloreactivity^12,13^. Thus, precise genome editing provides a generalizable strategy to enhance the safety and efficacy of T cell therapy^14^. However, the shift from random virus-based integration to precise CRISPR/Cas9-assisted insertion of CARs requires setting up a complete different T cell engineering platform.

The targeted integration of therapeutic genes needs the concomitant activity of three reagents, the CRISPR/Cas9 protein, a gRNA and a donor DNA template. Plasmids expressing gRNA and Cas9 are effective in cell lines but have limited efficiency in T cells^15^, so instead Cas9 mRNAs/proteins and synthesized gRNAs are delivered into T cells by electroporation^8–10,13^. After Cas9/gRNA makes site-specific cleavage, a HDR template is required for precise knock-in of the CAR gene. Using adeno-associated virus (AAV) to deliver HDR template can achieve high knock-in efficiency, generating ~20-45% edited T cell^8,11,16^. However, lengthy and expensive AAV production is a barrier to produce T cells for both research and clinical use^17^. AAV gene therapy also has potential genotoxicity^18^. Electroporated linear dsDNA has been explored as HDR template, producing ~7-20% edited T cells^9,10,14^. In detail, 4 μg linear dsDNA were needed for each 10^6^ T cells and the concentration has to reach 2 μg/μl for electroporation^9,10^, which can be achieved by pooling multiple PCR reactions with additional concentration steps. Even though producing linear dsDNA is easier and cheaper than AAV production, it is not as easy as extracting plasmid DNA from bacteria. Moreover, procedures to scale up PCR reactions and subsequent purification with Good Manufacturing Practices (GMP) to produce linear dsDNA haven’t been widely adopted for clinical applications yet. Meanwhile, large-scale GMP-grade plasmid manufacturing is commercially available and more economical than linear dsDNA manufacturing^19,20^. Thus, plasmid DNA, overcoming its current limitations, would be an ideal HDR template for both research and clinical use, provided that it could achieve comparable efficiency as linear dsDNA can.

Introducing two Cas9-cleavage sequences (CCSs) into HDR template plasmids has been shown to release the template by Cas9 and enhance knock-in efficiency in cell lines^21^. We first tested this strategy in 293T cells, introducing an N-terminal green fluorescent protein (GFP) fusion in the housekeeping gene *RAB11A*. There was a 75% increase of knock-in efficiency using template plasmids with two CCSs compared to those without CCSs, confirming previous findings^21^ (Supplementary Fig. 1a,b). Next we applied this strategy in primary T cells by co-delivery of Cas9/gRNA ribonucleoprotein (RNP) complex and CCSs-including plasmids (Fig. 1a). As our goal was to generate TRAC-CAR-T cells, we cloned the HDR template (~3.7 kb) for inserting a CD19-CAR at *TRAC* locus into the pUC57 vector (~2.7 kb) with two CCSs (Fig. 1a). Plasmids with two CCSs enhanced knock-in efficiency by 4.8 fold on average over plasmids without CCSs (n=2, Supplementary Fig. 2a,b). Since plasmid size affects the delivery efficiency by electroporation^22^, we deleted the *lacZ* module in the pUC57 backbone and generated a minimal vector (pMini, ~1.6 kb) to carry the HDR template for CAR knock-in (Fig. 1a). With this simple engineering, we can increase the knock-in efficiency by 8.6 fold on average, compared to pMini without CCSs (n=4, Fig. 1b,c). When T cells from different donors were sampled, CAR knock-in efficiency using pMini with two CCSs ranged from ~10% to 16.8% (n=11, Fig. 1d). In detail, when CD4 and CD8 T cells were analyzed separately, the two populations showed similar KI efficiency (Supplementary Fig. 3a,b). As plasmids with CCSs were also used for non-homologous end jointing (NHEJ)-mediated knock-in^23^, we sequenced the *TRAC* locus of edited cells and found all knock-in events were HDR-mediated (Fig. 1e,f). To compare the efficiency of our system with linear dsDNA-based non-viral method, we did a side-by-side knock-in test using linear dsDNA or plasmids with CCSs as HDR template while all other experimental parameters were kept the same^9,10^. Our results showed adding two CCSs to linear dsDNA significantly increased the efficiency (Supplementary Fig. 4a,b). We next assayed the CCSs-containing template plasmids for GFP fusion at *RAB11A* locus. The on-target knock-in efficiency can reach 8-9%, while off-target integration was minimal (Supplementary Fig. 5a-c).

**Figure 1.**
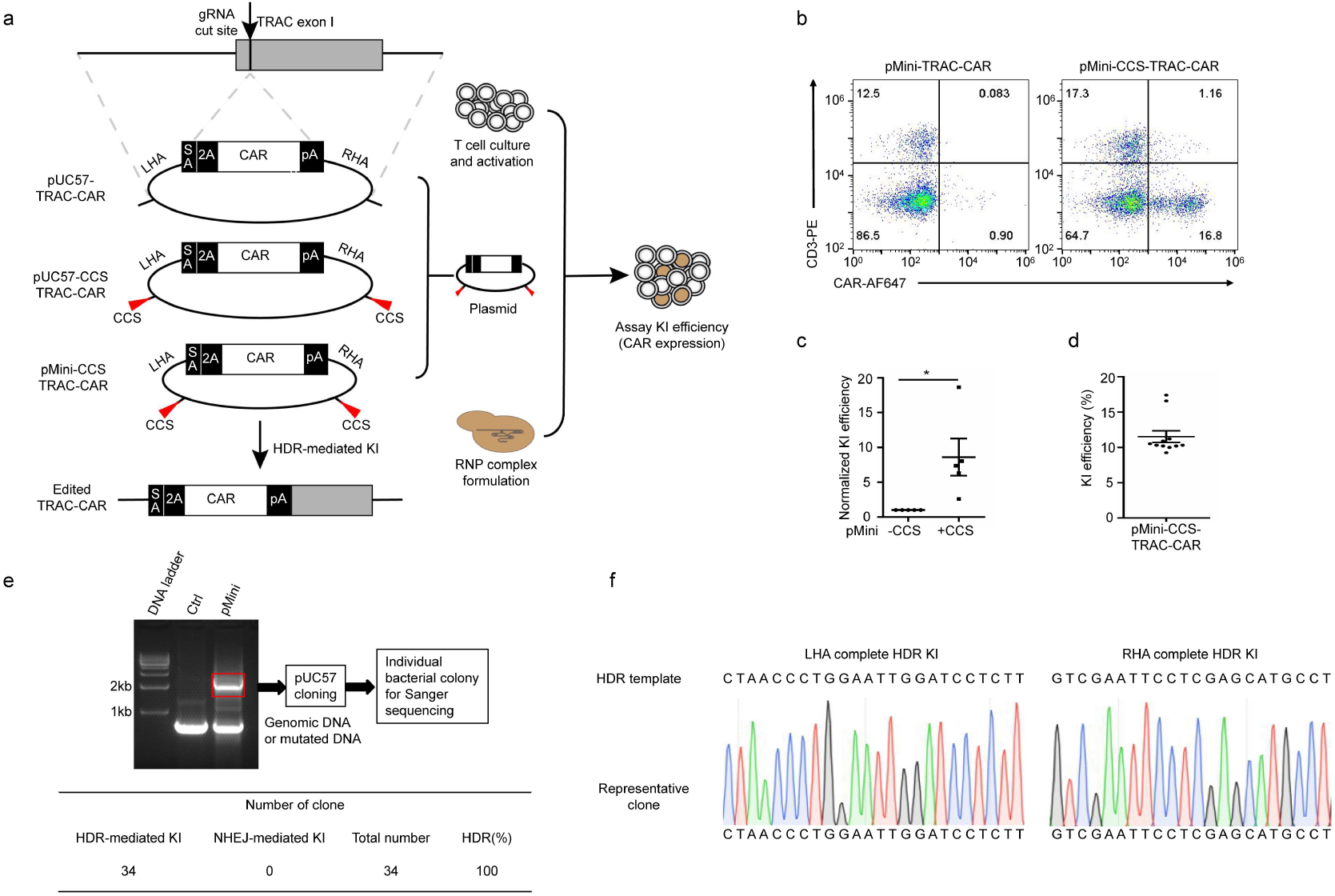
CRISPR/Cas9 mediated CAR knock-in (KI) with template plasmids in primary human T cell. **a.** Schematic of targeting a CAR to the *TRAC* locus with Cas9/gRNA RNP and different template plasmids. CCS, Cas9-cleavage sequence. **b.** Representative flow plots of CAR KI with indicated plasmids as template. **c.** Averaged fold increase of KI efficiency with pMini-CCS-TRAC-CAR compared to pMini-TRAC-CAR plasmids (n=5 donors). **d.** KI efficiency with pMini-CCS-TRAC-CAR averaged 11.7% (n=11 donors). **e.** The PCR product of 2-3kb size were cut off and cloned into pUC57 vector and individual bacterial colonies were picked for Sanger sequencing and summary of Sanger sequencing results. **f.** Alignment of sequencing result of a representative clone with expected HDR sequences at TRAC locus. Error bars represent S.E.M. *P < 0.05; **P <0.01; ***P < 0.001; ns, not significant (Student’s *t*-test).

Furthermore, our TRAC-CAR-T cells generated with template plasmids could be enriched from ~10% to more than 50% one week after stimulation with antigen presenting cells (APCs) (Fig. 2a). Such protocols to enrich CAR-T cells with APCs have been established for clinical use^24^. When different RNP and plasmid amounts were tested, no further enhancement was observed (Fig. 2b,c).Next we combined different strategies with the pMini vector to see if additional increase of knock-in efficiency could be achieved. As Cas9 with nucleus importing signals (NLSs) can help bring the template with CCSs into the nucleus, the number of CCSs in plasmids may affect the knock-in efficiency. When we systematically varied the number of CCSs in pMini from 0 to 3 (Fig. 2d), we observed the highest efficiency with 2 CCSs (Fig. 2e). Previous reports have suggested that homologous arm (HA) length can affect knock-in efficiency^21^. Therefore we constructed HDR templates with HA length varying from 100 to 800 bp. Our results showed that they all had similar knock-in efficiency (Fig. 2f). Moreover, *CCND1* gene was shown to further enhance the knock-in efficiency in cell lines^21^. When purified Cyclin D protein was electroporated together with Cas9/gRNA RNP and template plasmids, the knock-in efficiency was further enhanced by ~18% (Fig. 2g). We used MS-modified gRNA for all the experiments above as it was reported to be more stable than unmodified gRNA^25^. But it is more expensive, which could increase the manufacturing cost in clinical applications. Through side-by-side comparison, we found that MS-modified gRNA resulted in a 15% higher knock-in efficiency than unmodified gRNA (Fig. 2h). Thus we suggest using modified gRNA for T cell editing in research. But when editing large numbers of T cells in clinical settings, unmodified gRNA may be a cost-effective choice.

**Figure 2.**
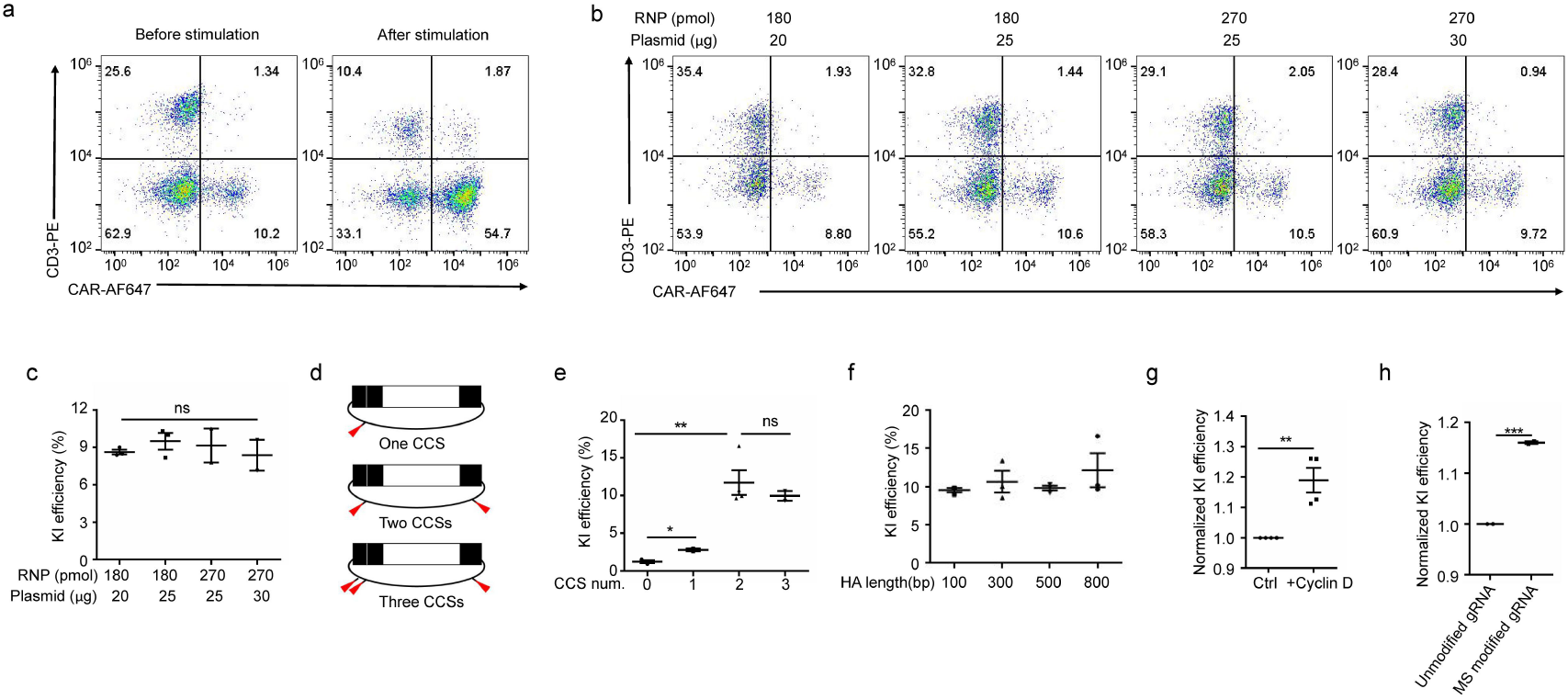
Optimization of CRISPR-Cas9 mediated CAR knock-in (KI) with template plasmids. **a.** Representative flow plots of CAR KI before or one week after stimulation with irradiated NIH/3T3-CD19 (n=3 donors). **b.** Representative flow plots of CAR KI with indicated amount of RNP and pMini-CCS-TRAC-CAR plasmid. **c.** Averaged KI efficiency with indicated amount of RNP and template plasmid (n=2 or 3 donors). **d.** Schematic of different CCS number located in pMini template plasmids. **e.** Averaged KI efficiency using pMini plasmids with indicated CCSs numbers (n=3 donors). **f.** Averaged KI efficiency with indicated HA length (n=3 donors). **g.** Averaged increase of KI efficiency with Cyclin D protein (n=4 donors). **h.** Comparison KI efficiency with normal guide RNA and MS guide RNA (n=2 donors). Error bars represent S.E.M. *P < 0.05; **P <0.01; ***P < 0.001; ns, not significant (Student’s *t*-test).

Next we assessed *in vitro* functions of these edited cells generated with plasmids (p-TRAC-CAR-T) and compared them vigorously with cells generated with AAV (AAV-TRAC-CAR-T). First, these p-TRAC-CAR-T cells kill CD19^+^ target cells as well as AAV-TRAC-CAR-T cells (Fig. 3a), which was reproducible in 3 donors. Second, both cells start to proliferate in the absence of antigen 3 days after electroporation, and continue to expand *in vitro*, providing sufficient cells for xenograft tumor eradication and clinical use (Fig. 3b). Third, we compared the long-term antigen-dependent proliferation in a weekly expansion assay with CD19^+^ APCs. The number of p-TRAC-CAR-T cells lagged behind that of AAV-TRAC-CAR-T cells in the first 4 weeks, but started to catch up and after 6 weeks ended up higher (Fig. 3c). The total 6-week expansion can generate as many as 1000 fold of cells. Fourth, we compared the genome-wide transcriptional profiles of p-TRAC-CAR-T and AAV-TRAC-CAR-T cells from two donors before and after antigen stimulation. Principal component analysis (PCA) demonstrated distinct clustering of CAR-T cells dependent on donor or stimulation status. However, p-TRAC-CAR-T and AAV-TRAC-CAR-T cells from the same donor before or after stimulation located closely, suggesting transcriptional similarity of these CAR-T cells (Fig. 3d). In addition, p-TRAC-CAR-T and AAV-TRAC-CAR-T cells showed similar expression profiles of genes differentially expressed between effector and naïve/memory T cells^26^ (Fig. 3e). A closer look of several key cytokines, exhaustion and activation markers confirmed the similarity (Supplementary Fig.6). Fifth, we used CyTOF mass cytometry to compare the phenotype (40 markers, Supplementary Table 1) of two types of CAR-T cells. t-Distributed Stochastic Neighbor Embedding (t-SNE) plot showed that p-TRAC-CAR-T and AAV-TRAC-CAR-T cells have comparable clustering patterns before or after antigen stimulation, suggesting the phenotypical similarity of these two types of cells (Fig. 3f). In detail, these differently made CAR-T cells showed similar levels of activation, differentiation and exhaustion markers (Supplementary Fig. 7).

**Figure 3.**
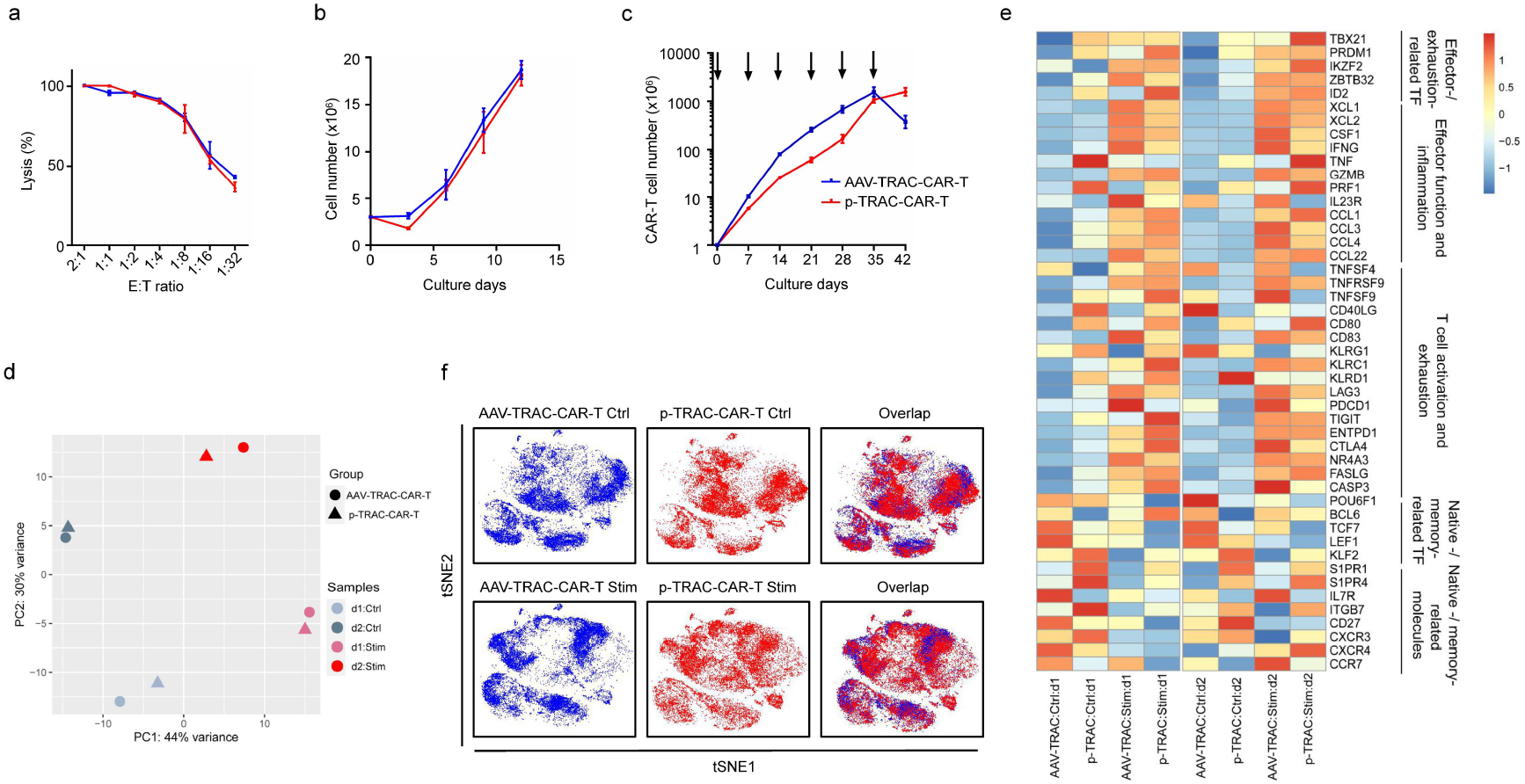
Functional comparison of edited TRAC-CAR T cells produced with plasmids or AAV as templates *in vitro*. **a.** Representative cytotoxic activity of CAR-T cells using an 18-h bioluminescence assay with FFL-Nalm6 as targets cells (n=3 donors performed in triplicates). **b.** Representative cell counts of total T cells on indicated days after electroporation (n=2 donors performed in triplicates). **c**. Representative cumulative cell counts of CAR-T cells upon weekly stimulation with irradiated NIH/3T3-CD19 cells (n=3 donors performed in replicates). **d.** Principal component analysis (PCA) of global transcriptional profiles (n = 2 donors). **e.** Heat map demonstrating the expression profiles of indicated genes for different CAR-T cells (n = 2 donors); TF, transcription factor. (d1, donor 1; d2, donor 2, Ctrl, before antigen stimulation; Stim, 24h after antigen stimulation) **f.** t-SNE analysis of p-TRAC-CAR-T and AAV-TRAC-CAR-T cells measured by 40 markers with or without stimulation by NIH/3T3-CD19 cells.

Last, we tested *in vivo* anti-tumour function of CAR-T cells in an acute lymphoblastic leukemia mouse model (Fig. 4a). where we injected the same low dose of AAV-TRAC-CAR-T cells and p-TRAC-CAR-T cells (1×10^5^). In these “stress test” conditions mice treated with different CAR-T cells showed no significant differences on the tumor burden (Fig. 4b, Supplementary Fig. 8) and long-term survival (Fig. 4c), suggesting that both CAR-T cells had equally potent anti-tumor activity. On day 16 after CAR-T cell injection, equivalent number of CAR-T cells accumulated in the bone marrow. On day 36, there were more p-TRAC-CAR-T cells than AAV-TRAC-CAR-T cells accumulated, suggesting longer persistence (Supplementary Fig. 9). The CAR we used above was a CD19-specific 1XX CAR developed for a range of clinical applications^26^. Since bispecific CARs have been shown to reduce relapse rate in clinical trials^27,28^, we tested whether a CD19 and CD22 bispecific CAR can be inserted at *TRAC* locus using the engineered plasmid pMiniZ (Fig. 4d), which was generated by replacing the ampicillin resistance gene in pMini with the zeocin resistance gene to further reduce the vector size. Even though the bispecific CAR (~2.2 kb) was about 50% larger than 1XX (~1.45 kb), the knock-in efficiency still reached 10.5% with the pMiniZ template (Fig. 4e), about 20% higher than that with pMini vector (Fig. 4f).

**Figure 4.**
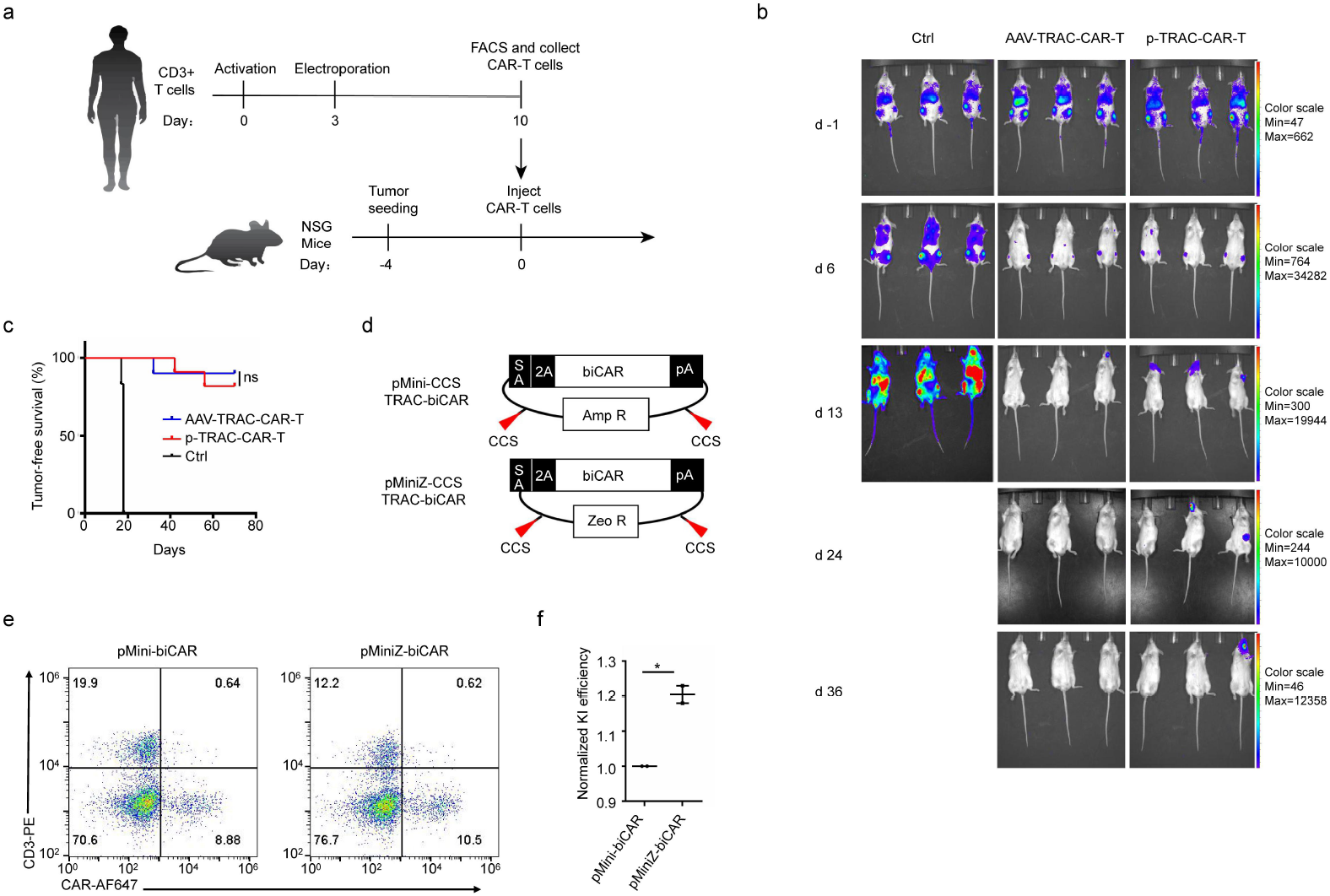
Functional comparison of edited TRAC-CAR T cells produced with plasmids or AAV as templates *in vivo*. **a.** Acute lymphoblastic leukemia tumour mouse xenograft model. NSG mice, non-obese diabetic (NOD)/severe combined immunodeficiency (SCID)/Il2rg-/- mice. **b.** Bioluminescent images of FFL-Nalm6-bearing mice treated with different CAR-T cells at indicated days (CAR-T cells were injected at day 0). **c.** Kaplan–Meier analysis of survival of mice treated with different CAR-T cells or untreated (Ctrl). (Ctrl: n=6, AAV-TRAC-CAR: n=10, p-TRAC-CAR: n=11). **d.** Schematic of bispecific CAR in pMiniZ or pMini template plasmids with CCSs. **e.** Representative flow plots of bispecific CAR KI with pMini-CCS-TRAC-biCAR and pMiniZ-CCS-TRAC-biCAR. **f.** Averaged KI efficiency using pMini-CCS-TRAC-biCAR and pMiniZ-CCS-TRAC-biCAR(n=2 donors). Error bars represent S.E.M. *P < 0.05; **P <0.01; ***P < 0.001; ns, not significant (Student’s t-test).

We have thus developed a plasmid-based non-viral method to rapidly and inexpensively knock-in genes in primary T cells. We reduced the size of the plasmid and flanked the HDR template with two CCSs to achieve efficient knock-in. Plasmid edited CAR-T cells were as potent as cells produced with AAV template, which was shown to have superior tumor eradication compared to virus-transduced CAR-T cells. This new method simplifies the gene targeting in human T cells and makes use of plasmid DNA that can easily be produced for clinical use. This approach will promote the use of precisely edited CAR-T cells for both research and clinical applications.

## Acknowledgement

We thank the support of Zhejiang Provincial Key Laboratory of Immunity and Inflammatory diseases. We thank Zhejiang University School of Medicine core facilities for their support. This research was funded by the National Natural Science Foundation of China grants 31971324 (J.S.) and 31971125 (C.Z.) and by Zhejiang Provincial Natural Science Foundation grant LR20H160003 (J.S.).

## Author contributions

R. J. and P. J. designed the study, performed experiments, analyzed and interpreted data. X. M., X. W., Y. D., K. S., L. Q., J. C, Y. H. and T. H. performed the experiments. J. L. and W. Y. analyzed data. J. J., W. C., X. Z. and X. G. interpreted data. C. Z., M. S. and J. S. designed the study, interpreted data and wrote the manuscript.

