## supplement information for "Cas9-cleavage sequences in minimal plasmids enhance non-viral genome targeting of CARs in primary human T cells"

### Methods

#### Cell lines and culture conditions

Nalm6 cells were transduced to express firefly luciferase (FFLuc)-green fluorescent protein (GFP), cultured in complete RPMI (Gibco) medium with 10% FBS (Vistech) and 1% Penicillin/Streptomycin (Gibco). NIH/3T3 cells were transduced to express human CD19 and cultured in DMEM (Gibco) medium with 10% FBS and 1% Penicillin/Streptomycin. 293T cells were cultured in DMEM medium with 10% FBS and 1% Penicillin/Streptomycin.

#### Isolation and expansion of human T cells

Peripheral blood mononuclear cells (PBMCs) were isolated by density gradient centrifugation from healthy volunteers' peripheral blood. Then T cells were purified using the Pan T Cell Isolation Kit (Miltenyi Biotec) and stimulated with CD3/CD28 T cell Activator Dynabeads (Thermo Fisher Scientific) with 1:1 cell to bead ratio and cultured in X-VIVO 15 serum-free hematopoietic cell medium (Lonza), supplemented with 10% FBS, 1% Penicillin/Streptomycin, 5 ng/ml interleukin-7 (IL-7) and 5 ng/ml interleukin-15 (IL-15) (Novoprotein). The medium was changed every 2-3 days, and cells were plated at $1\times10^6$/ml. Human T cells were stimulated for 48 hours, then debeaded for gene targeting experiments.

#### Plasmid construction

gRNA expression vectors were constructed by cloning 20 bp oligo nucleotide target sequences into pX330 (Addgene plasmid #42230) containing a human codon-optimized SpCas9 expression cassette and a human U6 promoter driving the expression of the gRNA.

The pMini and pMiniZ vectors were modified from a pUC57 vector. LacZ and multiple cloning sites were deleted from pUC57 to construct the pMini vector. Then the ampicillin resistance gene sequence was replaced with zeocin resistance gene sequence to make the pMiniZ vector.

To generate template plasmids harboring Cas9-cleavage sequences (CCSs) flanking the homology arms, the gRNA target sequence together with a PAM sequence (NGG) was included in both the forward and the reverse primers for cloning templates into the pUC57, pMini and pMiniZ vectors.

**Protein purification**

Cas9 from *S. pyogenes* with two NLS was subcloned from pX330 into a modified pET28b vector (Novagen), which contains an N-terminus 6× Histidine SUMO tag. Recombinant fusion protein was expressed in the *E. coli* strain Rosetta (Novagen). After purification with a HisTrap FF column (GE Healthcare), the his-sumo tag was cleaved by ULP1 and subsequently removed by a second step HisTrap FF column purification. Cas9 protein was further purified by a heparin column and a Superdex 200 increase column on AKTA Pure (GE Healthcare). The protein was concentrated to 10 mg/ml and stored in a buffer containing 20 mM HEPES pH 7.5, 200 mM NaCl, and 0.3 mM TCEP at -80 °C.

Human cDNA of *CCND1* was cloned into a modified pET28B vector, which contains an N-terminus 6× Histidine SUMO tag. Recombinant fusion protein was expressed in the *E. coli* strain Rosetta (Novagen). After purification with a HisTrap FF column (GE Healthcare), the his-sumo tag was cleaved by ULP1 and subsequently removed by a second step HisTrap FF column purification. Cyclin D protein was further purified with an ion-exchange column (Source Q) and Superdex 200 increase column on AKTA Pure (GE Healthcare). The protein was concentrated to 10 mg/ml and stored in a buffer containing 20 mM Tris–HCl pH 8.0, 200 mM NaCl, and 0.3 mM TCEP at -80 °C.

**RNP production**

The crRNAs used in this paper were C*A*G*GGUUCUGGAUAUCUGU for *TRAC* locus, G*G*T*AGTCGTACTCGTCGTC for *RAB11A* locus. The same tracrRNA AGCAUAGCAAGUUAAAAUAAGGCUAGUCCGUUAUCAACUUGAAAAAGU GGCACCGAGUCGGUGCU*U*U*U was used (* 2`-O-methyl 3`phosphorothioate, MS). RNPs were produced by complexing two components: gRNA and Cas9 protein. In brief, crRNAs and tracrRNAs were chemically synthesized with MS modifications and annealed (Genescript). The annealing was set up in a reaction containing 30 µl RNase free $H_2O$, 20 µl annealing buffer (5X, 50 mM Tris pH 8.0, 100mM NaCl), 25 μl crRNA (200 μM), 25 μl tracrRNA (200 μM), then heated at 95 ℃ for 5 minutes, gradually cooled to room temperature, aliquoted and lyophilized (Genescript). Lyophilized RNA was resuspended in RNase free water before use. 100 μM gRNA in 10 mM Tris pH 8.0, 20 mM NaCl were then mixed by 1:1.25 volume with 40 μM recombinant Cas9 (2:1 gRNA to Cas9 molar ratio) at 37 ℃ for 20 minutes to form 22 μM RNP complex. RNPs were then electroporated with template plasmids into T cells immediately after complexing.

**CAR-T cell production**

RNPs and template plasmids were electroporated into T cells 2 days after CD3/CD28 bead stimulation. Immediately before electroporation, debeaded T cells were centrifuged for 10 minutes, 1500 rpm and resuspended in BTXpress Electroporation Solution (BTX). For each reaction, $3\times10^6$ cells were mixed with 8.1 μl (22 μM, 180 pmol) RNPs and 7 μl (25 μg) template plasmids in a total volume of 100 μl and transferred to a 2 mm cuvette and electroporated with an AgilePluse system (BTX). Following electroporation cells were immediately transferred into culture medium and incubated at 37 ℃, 8% $CO_2$. 2 hours later, 5 ng/ml IL-7 and 5 ng/ml IL-15 were added. Then medium was changed every 2-3 days. For AAV-TRAC-CAR-T cells, 3.6 μl RNPs were incubated with $3\times10^6$ cells for electroporation. 30 minutes later, AAV virus (MOI=$1\times10^5$) (Vigene Biosciences, China) were added. 30 μg/ml Cyclin D protein was co-electroporated with RNPs and template plasmids into human T cells. 7 days after electroporation, cells were harvested for FACS analysis to determine the knock-in efficiency in each condition.

**Determination of HDR-mediated knock-in by PCR and Sanger sequencing**

Human T cells were harvested nine days after gene targeting for DNA extraction by FastPure cell/Tissue DNA Isolation Mini kit (Vazyme). The *TRAC* locus target sequences were amplified with Phanta HiFi DNA polymerase (Vazyme) with forward primer-F (GTTGTAAAACGACGGCCAGTGGTACCGCAGTATTATTAA GTAGCCC) and reverse primer-R (CTATGACCATGATTACGCCAAGCTT GTGGCAATGGATAAGGCCGAG). The bands in the size range of 2-3 kb were cut out and purified using a Gel Extraction Kit (Vazyme) and cloned into a pUC57 vector.

Individual bacterial colonies were picked for Sanger sequencing. Clones with high-quality sequencing data at both ends were aligned with expected HDR knock-in sequence by BLAST.

**Transfection of 293T cells**

For transfection of 293T cells, 25 kDa linear polyethyleneimine (PEI) (Polysciences) was used. 1 mg/ml PEI powder was dissolved in $H_2O$ and adjusted to pH 7.0 with HCl. For $1\times10^6$ cells in a 6-well plate, 1 μg pX330-RAB, 1 μg template plasmids and 6 μg PEI (1 mg/ml, PEI/DNA mass ratio is 3:1) were used.

**Antigen stimulation and proliferation assay**

CAR-T cell were co-cultured with irradiated 3T3-CD19 cells for weekly stimulation. $2.5\times10^5$ 3T3-CD19 cells were plated on 24-well tissue culture plates 12 h before addition of $5\times10^5$ CAR-T cells in X-VIVO 15 supplemented with FBS and cytokines. Total cells were counted and CAR expression was determined weekly by FACS. Subsequently, CAR-T cells were restimulated under the same conditions.

Cell staining

The following fluorophore-conjugated antibodies were used. From BD Biosciences: PE mouse anti-human CD8; BUV645 mouse anti-human CD4(Thermo Fisher Scientific); for CAR staining, an Alexa Fluor 647 AffiniPure F(ab' )$^2$ Fragment Goat Anti-Mouse IgG was used (Jackson ImmunoResearch). For TCR staining, a PE Mouse anti-Human CD3 was used (BD Bioscience). For cell counting, CountBright Absolute Counting Beads were added (Thermo Fisher Scientific) according to the manufacturer's instructions;For intracellular cytokine secretion assay, T cells were cocultured with 3T3-CD19 in the presence of the Golgi Plug protein transport inhibitor (BD Biosciences) for the 12 h. T cells were subsequently washed, stained for cell surface markers, fixed, and permeabilized according to the manufacturer's instructions using a Cytofix/Cytoperm Fixation/Permeabilization Solution Kit (BD Biosciences). Intracellular cytokine staining was performed with BV421 rat anti-human IL2, PE-Cy7 mouse anti-human TNF, FITC mouse anti-human IFN-γ (all BD Biosciences).

**Cytotoxicity assay**

Nine days after gene targeting, AAV-TRAC-CAR-T cells and p-TRAC-CAR-T cells were stimulated with irradiated 3T3-CD19 cells for a week and then collected for luciferase-based cytotoxicity assay using FFluc-GFP Naml6 as target cells. The effector (E) and target (T) cells were co-cultured in triplicates at indicated E/T ratios using black 96-well flat plates with $5\times10^4$ target cells in a total volume of 100 μl per well in RPMI medium. Target cell alone were plated at the same cell density to determine the maximal luciferase expression (relative light units (RLU)); 18 h later, 100 μl 150 ng/ml luciferase substrate (Goldbio) was directly added to each well. Emitted light was detected in a luminescence plate reader. Lysis was determined as (1-(RLUsample)/(RLUmax))×100.

**RNA extraction, sequencing and RNAseq analysis**

Seven days after gene targeting, AAV-TRAC-CAR-T cells and p-TRAC-CAR-T cells were stimulated with irradiated 3T3-CD19 for 24 h. RNA was extracted using TRIzol reagent (Thermo Fisher Scientific) followed by chloroform treatment and RNA precipitation by isopropyl alcohol. After RiboGreen RNA quantification and quality control using an Agilent 2100, samples were barcoded and run on a NovaSeq in a 150 base pair (bp)/150 bp paired-end run using the NovaSeq Reagent Kit (lllumina). The raw gene expression counts of each sample were annotated with the 'feature Counts' function in the R package 'Rsubread' based on the GRCh38 human genome reference. The integrated raw count matrix was then imported into the R package 'DESeq2' for downstream analyses, genes with more than 10 raw counts in at least 4 samples were kept and the filtered raw count matrix was transformed with the 'vst' (variance stabilizing transformation) method. These data were then applied for 2D projection with principal component analysis (PCA) and the scaled expression of selected genes were compared across samples to generate the heat map.

**CyTOF analysis**

Nine days after gene targeting, $3\times10^6$ T cells generated by AAV virus or template plasmids were stimulated with irradiated NIN/3T3-CD19 for 24 h, then harvested together with unstimulated T cells and sent for CyTOF analysis (PLT, Zhejiang, China). In brief, antibodies were either purchased pre-conjugated from Fluidigm

(DVS Sciences) or purchased purified and conjugated in-house using MaxPar X8 Polymer Kits (Fluidigm) according to the manufacturer's instructions. Cells for each sample were washed with protein-free PBS, stained with 0.25 μM Cell-ID Cisplatin-194Pt for 5 minutes at 4 °C, then incubated with blocking solution for 20 minutes at 4 °C, then stained for cell surface markers in staining media for 30 minutes at 4 °C. Cells were fixed and stained with DNA Intercalator-Ir (Fluidigm) overnight. Using the Foxp3/Transcription factor staining buffer set (eBiosciences) followed by intracellular staining for 30 minutes at room temperature, washed, and stored at 4 ℃ until acquisition. Cells were added with EQ Four Element Calibration Beads (Fluidigm) and analysed on a Helios instrument. All CyTOF fcs files were uploaded to PLT Biological Information Platform for subsequent analysis. The pre-proceed CyTOF fcs files (fcs 3.0 format) was obtained by standardizing and randomizing the original data. Also, all data has been debarcoded using a doublet filtering scheme with mass-tagged barcodes, and then manually gated to retain live, single, valid immune cells by using Flowjo Software. All cell events in each individual sample were pooled in this analysis. Data of all samples were analyzed by machine learning algorithm based on density estimation (Xshift), large-scale data clustering algorithm based on graph theory (PhenoGraph). Finally, the results were displayed by visualization methods such as TiSNE plot, heatmap, density plot and bar plot.

**Mouse systemic tumor model**

We used 6- to 12-week-old NOD/SCID/IL-2Rγ null mice (Biocytogen, China). All relevant animal-use guidelines and ethical regulations were followed. Mice were inoculated with $0.5\times10^6$ FFluc-GFP Naml6 cells by tail vein injection, followed by $1\times10^5$ CAR-T cells injected 4 days later. Bioluminescence imaging used the IVIS Imaging System (PerkinElmer) with Living Image software (PerkinElmer) for acquisition of imaging datasets. At each indicated time points, 3 mice from each group were imaged and measured for average tumor burden.

**Isolation of cells from bone marrow and spleen**

Mice were euthanized with $CO_2$ at day 16 and day 36 after CAR-T cell injection. Bone marrow was harvested from freshly isolated femurs and tibiae. After removal of connective tissues and muscles, bones and spleens were crushed in 5ml PBS-EDTA. Single-cell suspensions were made by pipetting and passing supernatant from bone marrow and spleen through a 40 μm filter (BD Falcon). Remaining RBCs were lysed with ACK buffer.

**Statistical analysis**

All statistical analyses were performed using the Prism 5 (GraphPad) software. No statistical methods were used to predetermine sample size. Statistical comparisons between two groups were determined by two-tailed paired Student's *t*-tests for matched samples. For *in vivo* experiments, the overall survival was depicted by a Kaplan–Meier curve and the log-rank test was used to compare survival differences between the groups. P values $< 0.05$ were considered to be statistically significant. The statistical test used for each figure was described in the corresponding figure legend.

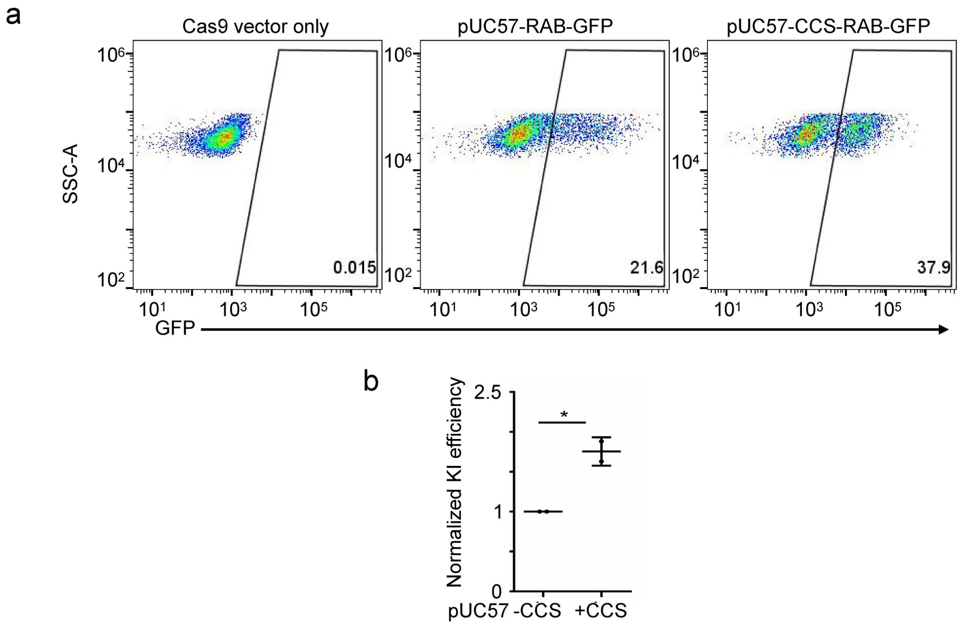


**Supplementary Fig. 1** Including two CCSs in the template plasmid resulted in enhanced gene knock-in (KI) efficiency in 293T cells. **a.** Representative flow plots of GFP KI at *RAB11A* locus with or without CCSs in template plasmids 7 days after transfection. **b.** Averaged fold increase of normalized KI efficiency with pUC57-CCS-RAB-GFP compared to pUC57-RAB-GFP plasmids (n=2 replicates). Error bars represent S.E.M. *P < 0.05 (Student's *t*-test).

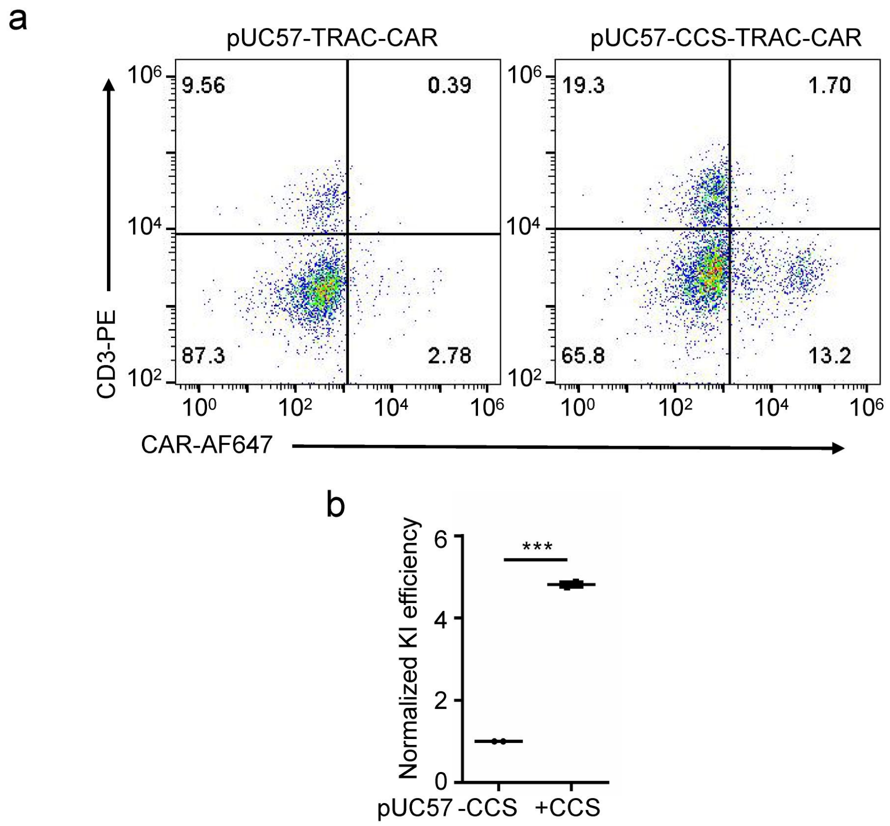


**Supplementary Fig. 2** Including two CCSs in the template plasmid pUC57 resulted in enhanced gene KI efficiency in primary human T cells. **a.** Representative flow plots of CAR KI with indicated template plasmids 7 days after electroporation. **b.** Averaged fold increase of normalized KI efficiency with pUC57-CCS-TRAC-CAR compared to pUC57-TRAC-CAR plasmids (n=2 donors). Error bars represent S.E.M. ***$P < 0.001$ (Student's *t*-test).

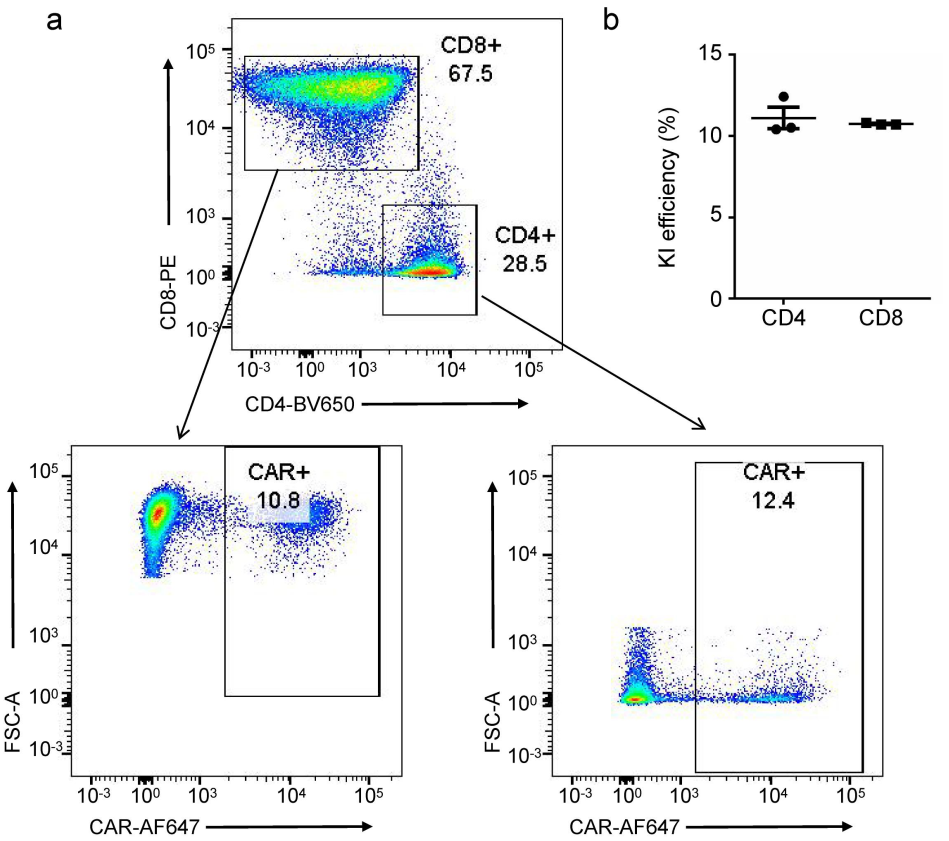


**Supplementary Fig. 3** CAR KI efficiency in primary human CD4 and CD8 T cells. **a.** Representative flow plots **of** the CAR KI efficiency in CD4 and CD8 T cell populations. **b.** Average KI efficiency in CD4 and CD8 cell populations (n=3 donors).

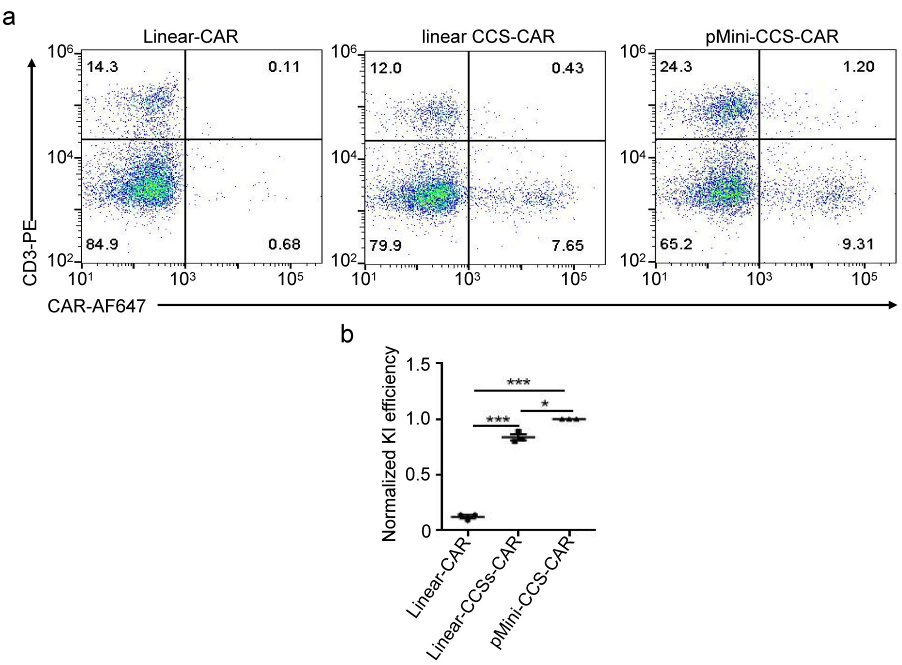


**Supplementary Fig. 4** pMini-CCS-TRAC-CAR as template has higher KI efficiency than linear-CCS-TRAC-CAR and linear-TRAC-CAR in primary human T cells. **a.** Representative flow plots of CAR KI with indicated templates (n=3 donors). **b.** Averaged fold change of KI efficiency with indicated templates normalized to pMini-CCS-TRAC-CAR as template (n=3 donors). Error bars represent S.E.M. *P < 0.05; **P <0.01; ***P < 0.001 ; (Student's t-test).

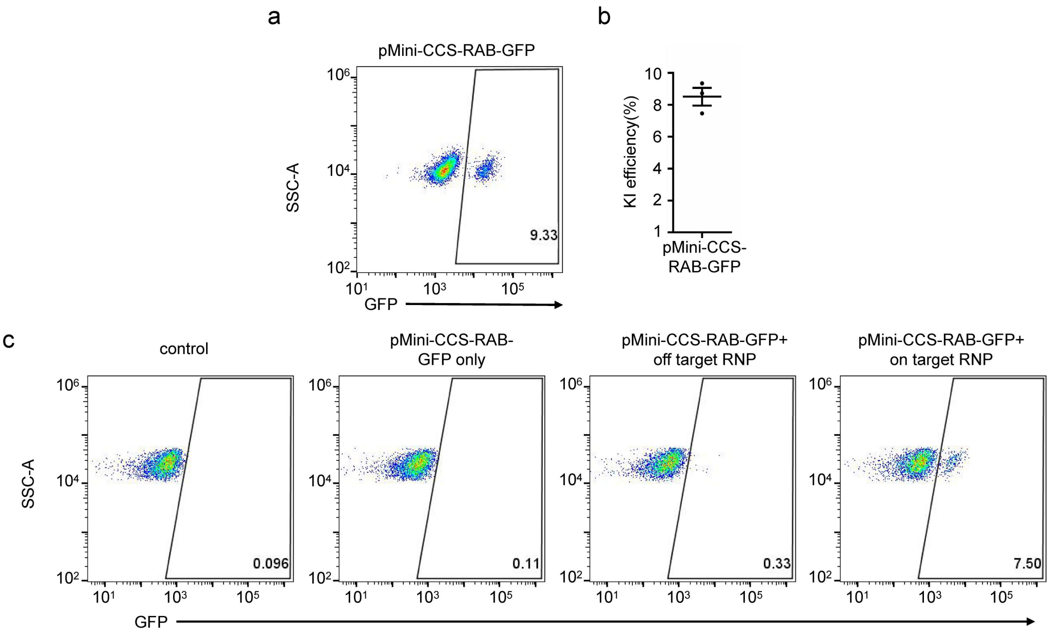


**Supplementary Fig. 5** The KI of GFP fusion at *RAB11A* locus in primary human T cells. **a.** Representative flow plots of GFP KI efficiency 7 days after electroporation. **b.** Averaged KI efficiency at *RAB11A* locus. **c.** For possible off-target RNP, unintended non-homologous integration with pMini-CCS-RAB-GFP template was minimal.

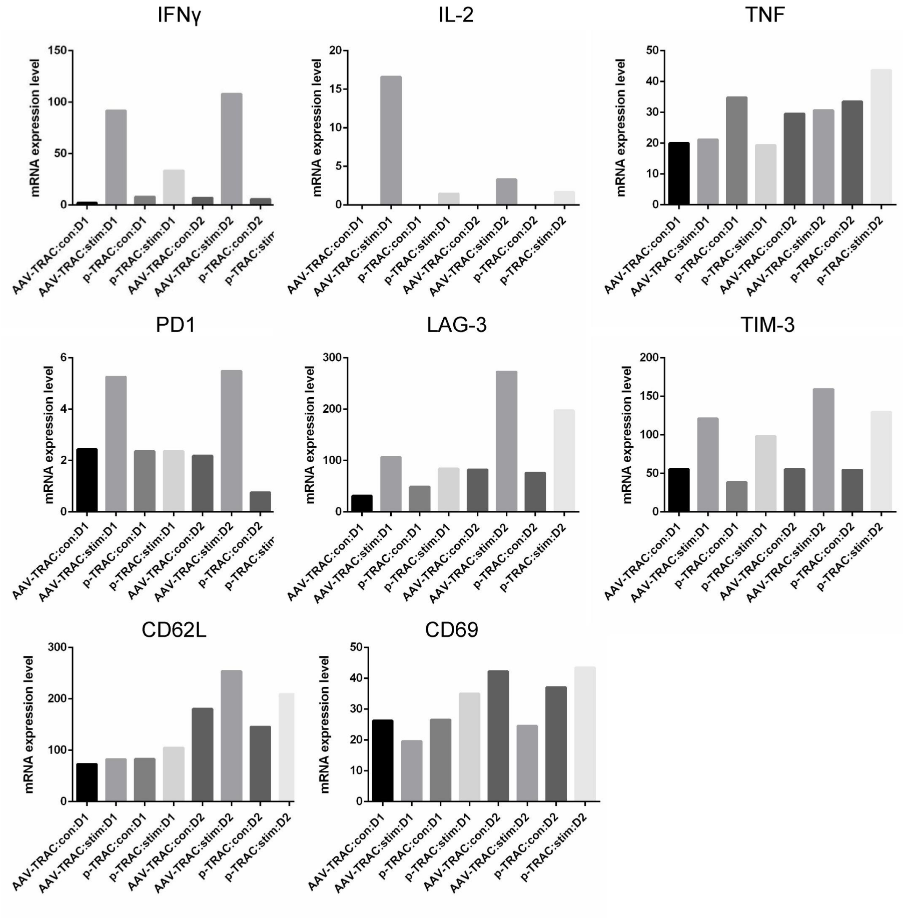


**Supplementary Fig. 6** The transcription level of key cytokines (IL2, IFN $\gamma$ , TNF), exhaustion markers (PD-1, LAG-3, TIM-3), activation marker (CD69) and memory marker (CD62L) in indicated CAR-T cells extracted from RNAseq data. (d1, donor 1; d2, donor 2, con, before antigen stimulation; stim, 24h after antigen).

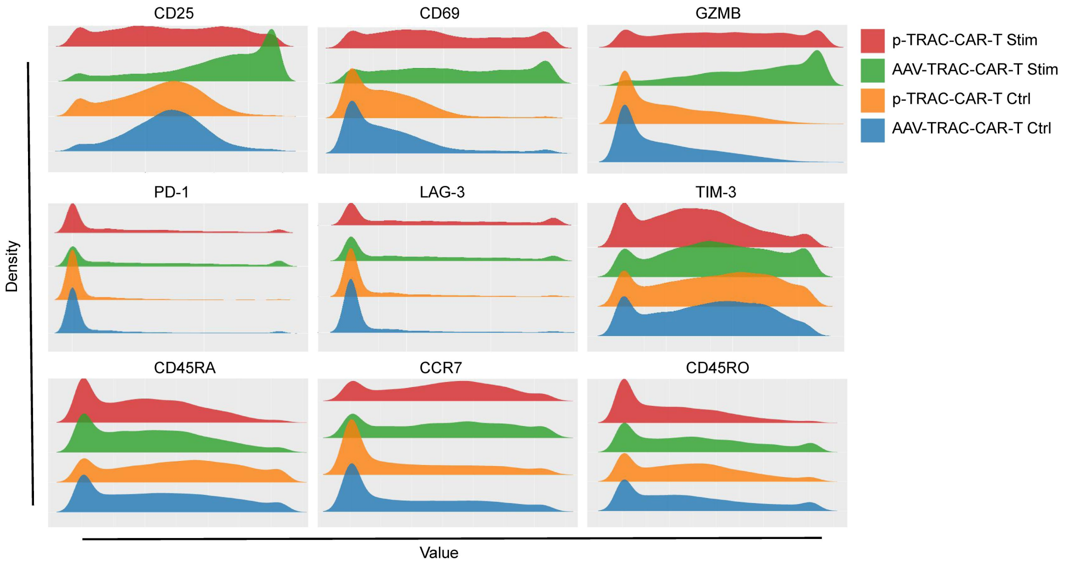


**Supplementary Fig. 7** The expression level of T cell activation markers (CD25, CD69), Granzyme B, exhaustion markers (PD-1, LAG-3, TIM3) and differentiation markers (CD45RA, CCR7, CD45RO) in indicated CAR-T cells.（Con, before antigen stimulation; Stim, 24 h after antigen stimulation).

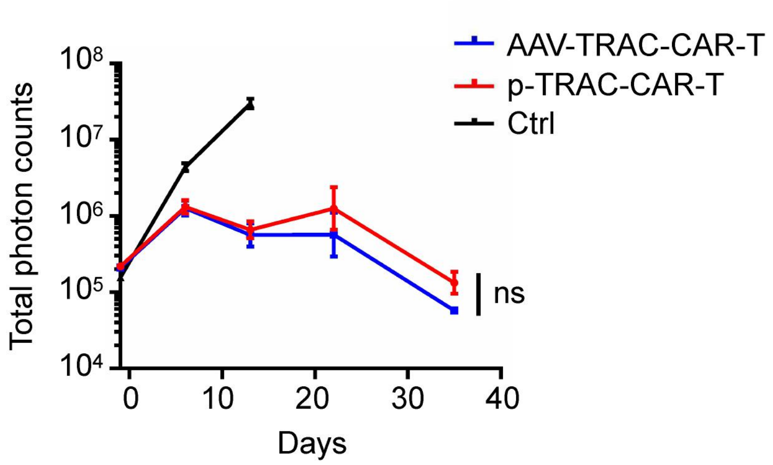


**Supplementary Fig. 8** Averaged tumor burden of FFL-NALM6-bearing mice treated with indicated CAR-T cells at different time points (CAR-T cells were injected at day 0). Error bars represent S.E.M. ns, not significant (n=6, Student's *t*-test).

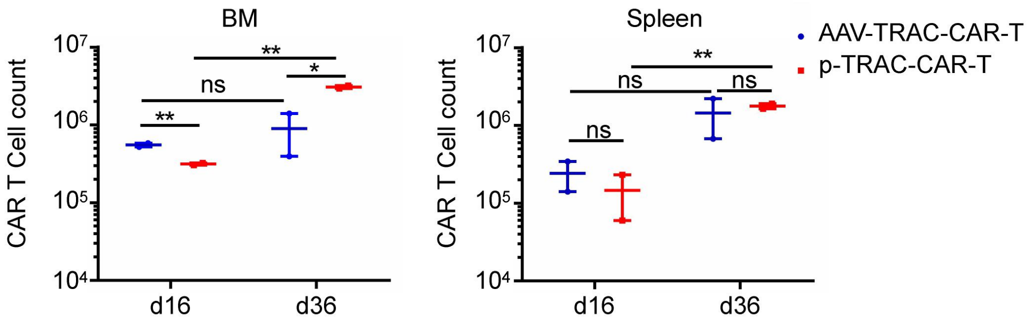


**Supplementary Fig. 9** FFL-Nalm6-bearing mice were treated with $1\times 10^5$ indicated CAR-T cells. CAR-T cells isolated from bone marrow and spleen were counted 16 and 36 days after administration. Error bars represent S.E.M. ns, not significant; *$P < 0.05$; **$P < 0.01$(n=2, Student's *t*-test).

| CyTOF detected markers | | | |
|---|---|---|---|
| CD3 | CD7 | CD28 | CD127 |
| CD4 | CD38 | CD152_CTLA4 | Gata3 |
| CD8a | CD223_Lag3 | FoxP3 | CD366_Tim3 |
| CD19_Pro | CD161 | CD137_41BB | GZMB |
| OX40_CD134 | Ki67 | RORγ | CD279_PD1 |
| CD95_FAS | CD45RA | CD278_ICOS | CCR10 |
| CD193_CCR3 | CD194_CCR4 | CD314_NKG2D | HLA_DR |
| CD183_CXCR3 | CD27 | CD45RO | CD196_CCR6 |
| CD69 | CD197_CCR7 | T-bet | γδTCR |
| CD25 | CD357_GITR | CD185_CXCR5 | |

**Supplementary Tab. 1** The list of markers used by CyTOF mass cytometry.
